# Neuroprotective, antioxidant and antiproliferative activity of grapefruit IntegroPectin on SH-SY5Y cells

**DOI:** 10.1101/2021.07.21.453202

**Authors:** Domenico Nuzzo, Miriana Scordino, Antonino Scurria, Costanza Giardina, Francesco Giordano, Francesco Meneguzzo, Giuseppa Mudò, Mario Pagliaro, Pasquale Picone, Alessandro Attanzio, Stefania Raimondo, Rosaria Ciriminna, Valentina Di Liberto

## Abstract

Tested *in vitro* on SH-SY5Y neuroblastoma cells, grapefruit IntegroPectin is a powerful neuroprotective, antioxidant and antiproliferative agent. The strong antioxidant properties of grapefruit IntegroPectin, and its ability to preserve mitochondrial membrane potential and morphology, severely impaired in neurodegenerative disorders, make this new biopolymer highly soluble in water an attractive therapeutic agent for oxidative stress-associated brain disorders. Similarly, the ability of this new citrus pectin rich in naringin, linalool, linalool oxide and limonene adsorbed at the outer surface to inhibit cell proliferation or even kill, at high doses, neoplastic cells, coupled to its excellent health and safety profile, opens up new therapeutic strategies in cancer research. In order to take full advantage of its vast therapeutic and preventive potential, detailed studies of molecular mechanism involved in the antiproliferative and neuroprotective of IntegroPectin are urgently needed.

## 1. Introduction

Globally contributing 16.5% of deaths from all causes and 11.6% of global disability-adjusted life-years in year 2016, neurological disorders (NDs) are the second leading group cause of deaths in the world [1]. In recent years NDs have increased significantly due to the ageing of the population, malnutrition, various forms of environmental pollution, lifestyle, diet [2], viral infections and other environmental and social factors [3]. Causing cell damage, oxidative stress is one of the main mechanisms involved in NDs, since it alters numerous cellular processes such as mitochondrial homeostasis [4], DNA repair and cell signalling propagating cell damage and leading to incurable neurodegenerative diseases [5].

To date, no effective synthetic or natural drugs are available for preventing o treating NDs such as Alzheimer’s, Parkinson’s disease, and amyotrophic lateral sclerosis. Hence, plentiful research has been devoted to search bioactive natural compounds to be used as neuroprotective and neuroregenerative agents. For example, algae-derived antioxidant molecules such as phycocyanins have been successfully used to inhibit cellular oxidative stress, mitochondrial dysfunction and apoptosis, and increase neuronal viability in an *in vitro* model of Alzheimer’s disease [6,7]. In general, antioxidant molecules from naturally derived sources exhibit high bioavailability and significantly greater efficacy than synthetic antioxidants [8]. A regular and correct intake of natural antioxidants plays an important role in the prevention and control of NDs, with several phytochemical complexes with antioxidant activity having been identified as potential ND therapeutic agents [9].

In this context, pectin, the ubiquitous plant heteropolysaccharide is emerging as a “universal medicine” thanks to its immunomodulating properties, including anti-inflammatory activity [10]. Comprised of structurally distinct homogalacturonan (HG, accounting for 65%), rhamnogalacturonan (RG-I, accounting for 20-35%), and rhamnogalacturonan (RG-II, accounting for 2–10%) regions, pectin is a hydrocolloid abundant in the cell wall where it acts as glue ensuring plant cell adhesion [11].

The biopolymer is a natural component of all omnivorous diet and an important soluble dietary fiber promoting numerous beneficial effects on intestinal microbiota, preventing several inflammatory conditions [12]. Owing to its excellent palatability, ability to provide texture at low concentration, and excellent health and safety profile, pectin is the most valued natural hydrocolloid used by the food industry, with demand increasing at fast pace since more than a decade [13]. From new encapsulant in pharmaceutical pills through use of enzymatically degraded pectin as nutraceutical ingredient, furthermore, a number of new applications of pectin have lately emerged which also contribute to its rapidly expanding demand [14]. The industrial production process, no longer based as in the early 1900s on cooking apple pomace, is based on hydrolysis of dried or wet citrus (lemon, orange and lime) peels and apple pomace with mineral acid in hot water followed by precipitation with isopropyl alcohol [15]. The process significantly degrades the structure of natural pectin, especially hydrolyzing the side chains of the RG-I “hairy” region [16]. Accounting for 20-35% of pectin, the latter region is composed of a backbone of repeating galacturonic acid and rhamnose disaccharide with neutral sugar side chains attached to the *O*-4 position and sometimes the *O*-3 position of rhamnose backbone units. The neutral sugar side chains consisting of arabinose and galactose with variable linking types and chain lengths have demonstrated biological function, strongly binding and inhibiting galectin-3 and thus leading to anti-apoptotic activity [17].

From ultrasound-assisted and microwave-assisted extraction numerous non-thermal and thermal methods to extract RG-I enriched pectins have been lately developed [16]. Among the latter, hydrodynamic cavitation has been lately first applied to waste orange peel [18] as well as to lemon and grapefruit [19] biowaste resulting from *Citrus* fruit juice extraction. Rich in flavonoids [20] and terpenes [21], the IntegroPectin resulting from freeze-drying of the aqueous hydrocavitated extract has exceptionally high antioxidant activity and lacks cytotoxicity even at high concentration [19]. Furthermore, lemon and grapefruit IntegroPectin show significant antibacterial activity [22]. Subsequent detailed investigation has shown high and broad-scope antimicrobial activity for both newly obtained pectins, particularly for grapefruit IntegroPectin which is a bactericidal agent at low concentrations for both Gram-positive and Gram-negative pathogenic bacteria [23]. Recently, the powerful *in vitro* mitoprotective and neuroprotective activity of lemon IntegroPectin on neuronal SH-SY5Y human cells treated with concentrated (0.2 M) aqueous H_2_O_2_, a strong oxidizer involved in the cellular mechanisms leading to neurodegenerative pathologies, has been reported [24]. In this study, along with high neuroprotective and antioxidant effect of grapefruit IntegroPectin on the same neuronal model cells, we report the discovery of powerful antiproliferative activity.

## 2. Results

### 2.1. Effect on SH-SY5Y cell viability

We first evaluated the effect of different concentrations of grapefruit IntegroPectin on cell viability of SH-SY5Y cells. As shown in Figure 1a, displaying the outcomes of the MTT (3-(4,5-dimethylthiazol-2-yl)-2,5-diphenyltetrazolium bromide) colorimetric assay, 24 h treatment with the new citrus pectin caused a dose-dependent reduction in cell viability, which became significant for doses exceeding 0.1 mg/ml. At 10 mg/ml dose, only a few cells remained viable. Similar results were obtained for treatment prolonged up to 48 h, and in human lung carcinoma cells H292 (data not shown). However, while the dose of 10 mg/ml produced clear cytotoxic effects, as evidenced by morphological analysis (Figure 1b), the 1 mg/ml dose was not associated to alteration of cell morphology (Figure 1b) and to significant increase in cell death, as shown by the cytotoxicity (CellTox) assay (Figure 1c) and by the healthy morphology of cell nuclei counterstained with DAPI (4′,6-diamidino-2-phenylindole) fluorescent dye (Figure 1d).

**Figure 1.**
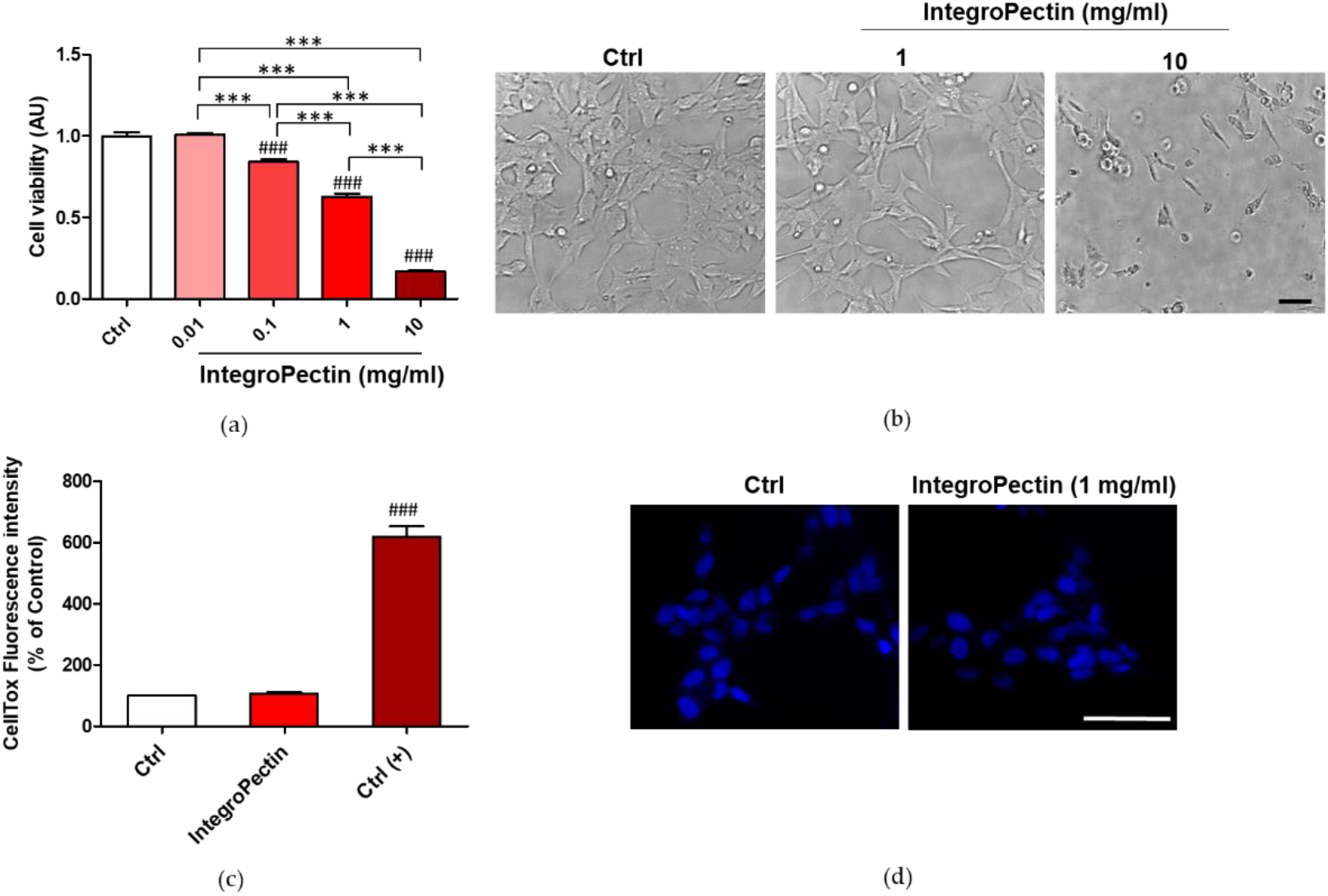
Dose dependent effects of grapefruit IntegroPectin on neuronal cell viability. (a) Cell viability of Integropectin treatment (24 h) in dose dependent experiment; (b) Representative morphological images of untreated cells (Ctrl) or cells treated with different doses (1, 10 mg/ml) of IntegroPectin (24 h); (c) Cytotoxicity associated to IntegroPectin treatment (1 mg/ml, 24h), evaluated by fluorescence developed by CellTox Green Cytotoxicity Assay. Positive control Ctrl (+) is represented by cell exposure to lysis buffer; (d) Microscopy inspection of DAPI-labeled nuclei in Ctrl cells or treated with IntegroPectin (24 h). ### p < 0.001 as compared to control (Ctrl) group; *** p < 0.001. Scale bar 50 μm.

Therefore, in order to understand the results of the MTT test and the absence of cell death at the dose of 1 mg/ml, we next analyzed the eventual cytostatic effects.

### 2.2. Cytostatic effect on SH-SY5Y cells

The distribution of cells in the different phases of the cell cycle, analyzed by cytofluorimetry analysis of cellular DNA content following cell staining with propidium iodide (PI), shed insight on the cytostatic effect of the new grapefruit pectin on neuronal model cells. As shown in Figure 2, treatment with grapefruit IntegroPectin produces a cell cycle arrest exactly at the G_2_/M phase. We briefly remind that the growth 2 phase (G_2_ phase) is the third subphase of interphase in the cell cycle directly preceding mitosis, and that cell cycle arrest at the G_2_/M phase indicates that the damage of intracellular DNA is difficult to repair [25]. The G_2_/M-phase checkpoint usually prevents cells with damaged DNA from undergoing mitosis by the inhibition of the mitotic complex CDK1-cyclin B and activation of the apoptosis cascade [26]. However, we did not detect any increase in cell death when treatment was prolonged up to 5 days (data not shown).

**Figure 2.**
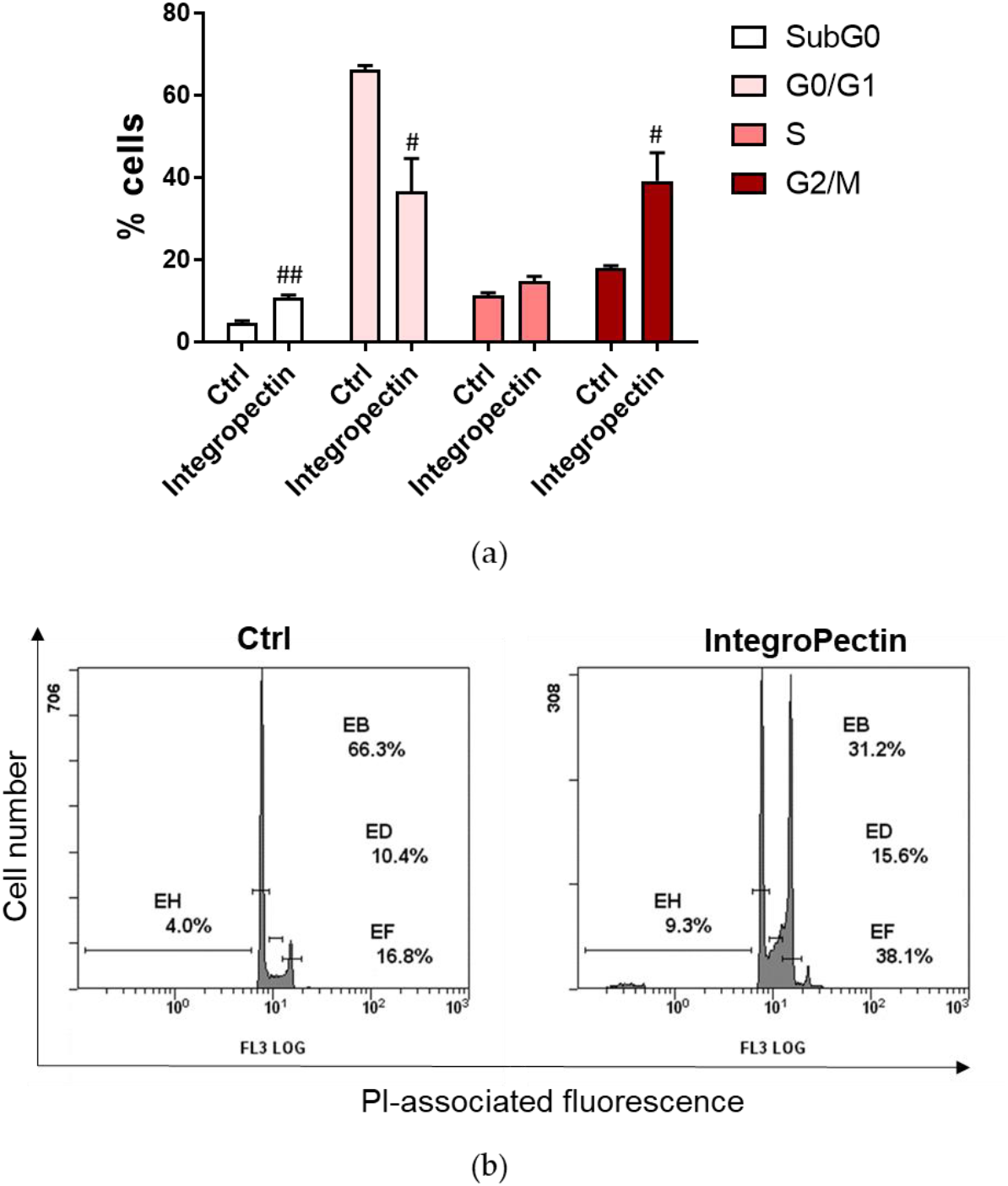
Effect of grapefruit IntegroPectin on the cell cycle distribution of SH-SY-5Y cells. Percentage (%) of cell distribution of untreated (Ctrl) cells and cells treated for 24 h with IntegroPectin (1 mg/ml) in different phases of the cell cycle, assessed by flow cytometry analysis after propidium iodide (PI) staining; (b) Representative images. # p < 0.05, ## p < 0.01 as compared to control (Ctrl) group

We thus assessed the potential antioxidant and neuroprotective properties of the new citrus pectin in cells exposed to concentrated (0.2 M) aqueous H_2_O_2_.

### 2.3 Neuroprotective effect on SH-SY5Y cells

Pretreatment (Figure 3a) with grapefruit IntegroPectin at a dose of 1 mg/ml, *per se* cytostatic, was able to significantly counteract cell death induced by the treatment with 0.2 M H_2_O_2_ (Figure 3b). The same treatment was also able to recover cell morphology and cell body area, impaired by H_2_O_2_ treatment (Figures 3c and 3d). Furthermore, Figures 3c and 3e show that pretreatment with the newly sourced citrus pectin was able to reduce the number of cell debris, indicative of cell protection. Similar results were obtained when the new grapefruit IntegroPectin was applied along with aqueous H_2_O_2_ (data not shown).

**Figure 3.**
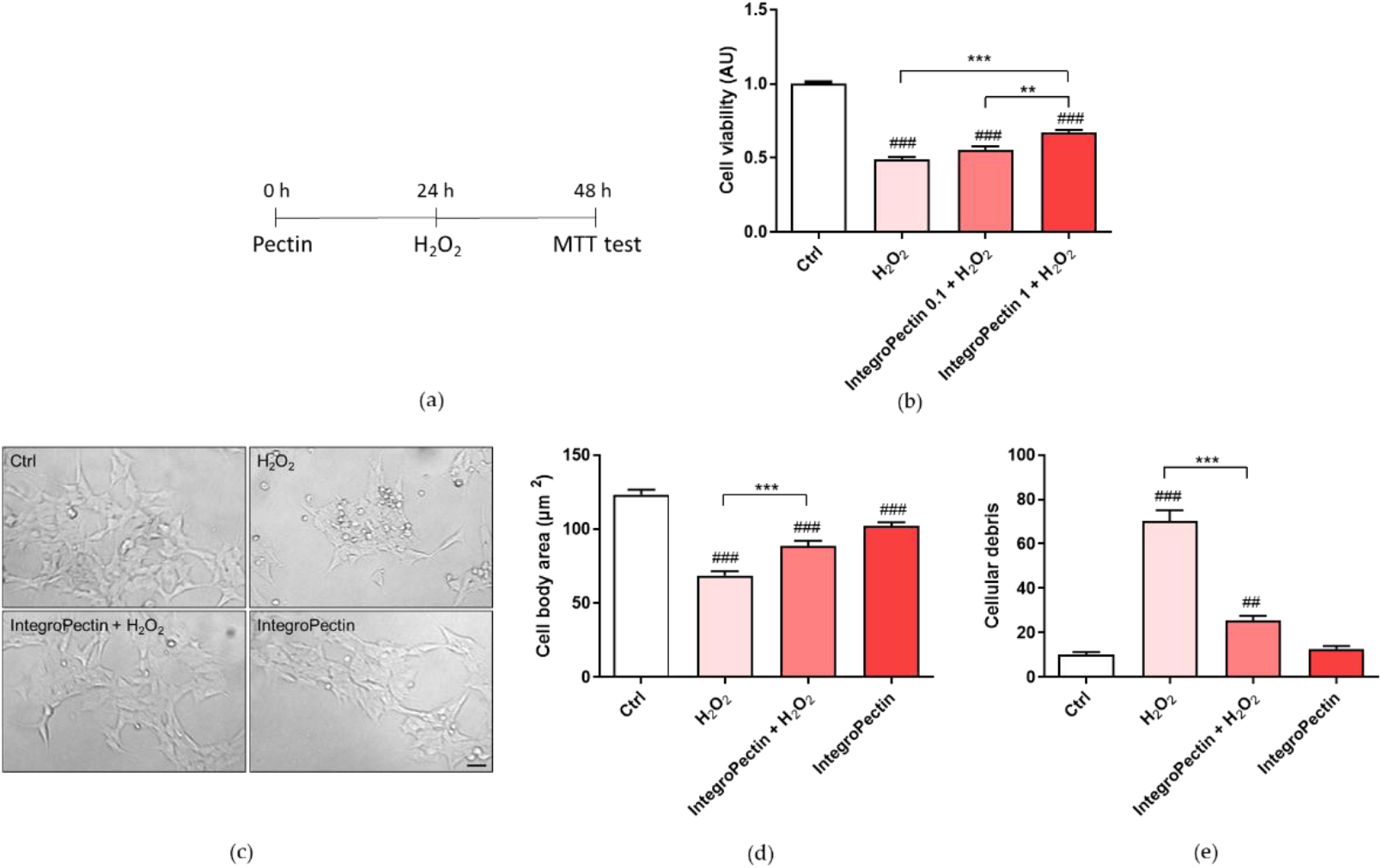
Effect of grapefruit IntegroPectin on cell viability and morphology of SH-SY5Y cells impaired by H_2_O_2_ treatment. (a) Scheme of cell pretreatment with IntegroPectin; (b) Histogram showing cell viability of untreated cells (Ctrl) or treated with IntegroPectin or with H_2_O_2_ alone or in combination with IntegroPectin; (c) Representative bright field morphological images of untreated cells (Ctrl) or treated with IntegroPectin or with H_2_O_2_ alone or in combination with IntegroPectin; (d) Cell body area histogram of untreated cells (Ctrl) or treated with IntegroPectin or with H_2_O_2_ alone or in combination with IntegroPectin; (e) Number of cells debris histogram of untreated cells (Ctrl) or treated with IntegroPectin or with H_2_O_2_ alone or in combination with IntegroPectin. Scale bar 50 μm. Tukey test: ## p < 0.01, ### p < 0.001 as compared to control (Ctrl) group; ** p < 0.01, *** p < 0.001.

### 2.4 Antioxidative effect on SH-SY5Y cells

The impact of grapefruit IntegroPectin on H_2_O_2_-induced oxidative stress was assessed measuring reactive oxygen species (ROS) production by the dichlorofluorescein diacetate (DCFH-DA) fluorescence intensity assay, a widely used probe for detecting oxidative stress and intracellular reactive species [27]. Fluorescence microscope inspection (Figure 4a) and fluorescence intensity measurement (Figure 4b) showed that treatment of neuronal cells with grapefruit IntegroPectin almost completely counteracted ROS formation driven by exposure to concentrated H_2_O_2_. The kinetics of ROS production after exposure of SH-SY5Y cells to H_2_O_2_ shows that treatment with IntegroPectin is indeed quickly and highly effective in lowering and delaying ROS production due to hydrogen peroxide addition (Figure 4c).

**Figure 4.**
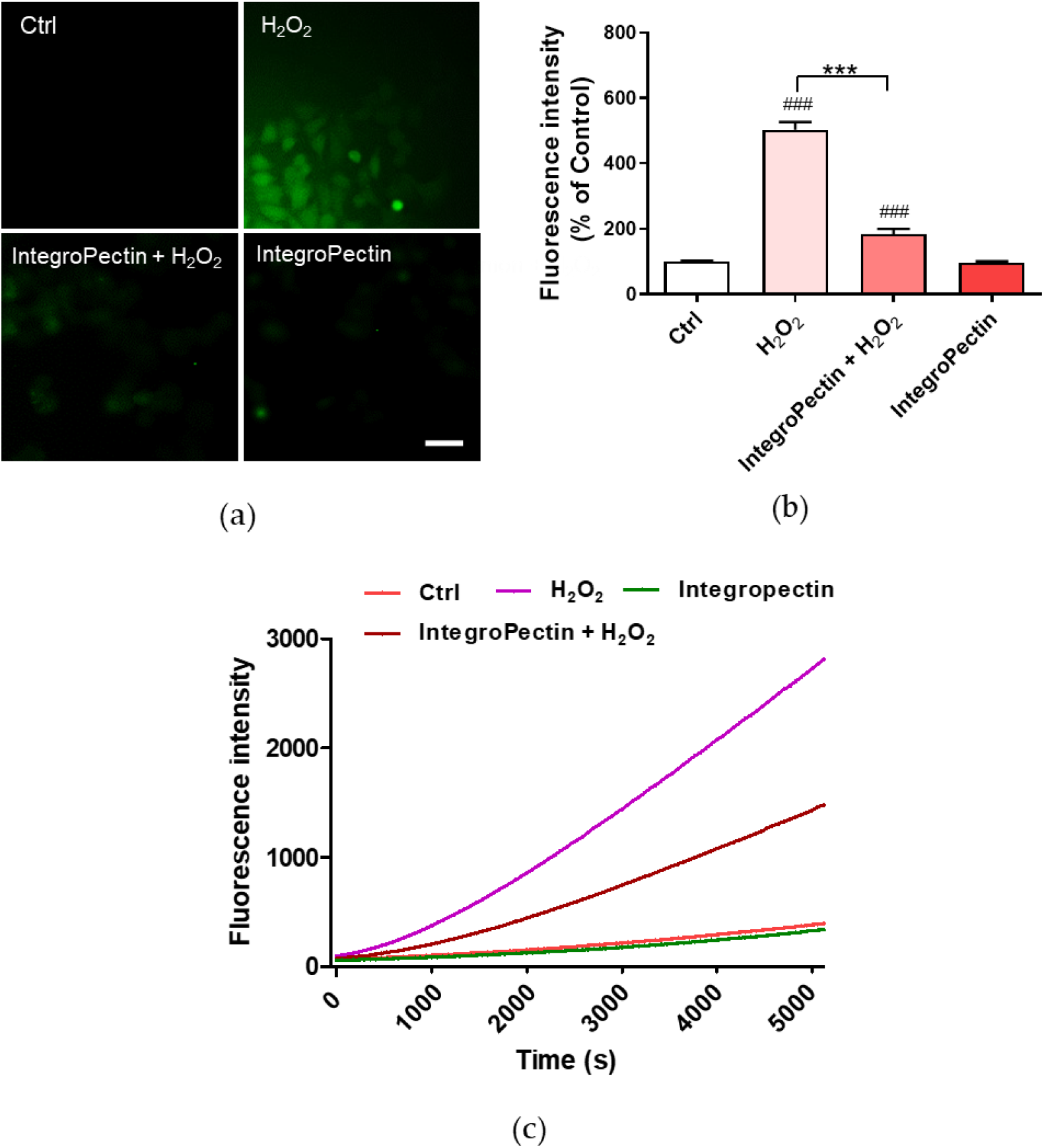
Effects of IntegroPectin on ROS production driven by exposure to aqueous H_2_O_2_. (a) DCFH-DA fluorescence microscopy images of untreated cells (Ctrl) or treated with IntegroPectin or with H_2_O_2_ alone or in combination with IntegroPectin; (b) Histogram of fluorescence intensity of untreated cells (Ctrl) or treated with IntegroPectin or with H_2_O_2_ alone or in combination with IntegroPectin measured by DCFH-DA fluorescence assay; (c) Oxidation kinetics of untreated cells (Ctrl) or treated with IntegroPectin or with H_2_O_2_ alone or in combination with IntegroPectin, monitored by DCFH-DA fluorescence assay. Scale bar: 50 μm. Tukey test: ### p < 0.001 as compared to control (Ctrl) group, *** p < 0.001.

### 2.5 Mitoprotective effect on SH-SY5Y cells

Variations in the physiological mitochondrial membrane potential, an indicator of cells’ health and functional status in response to oxidative stress, were measured as changes in the accumulation of JC-1 cyanine dye red and green fluorescence signals in the cells. When excited at 488 nm, JC-1 monomers emit green fluorescence with a maximum at 530 nm (green), whereas J-aggregates emit orange-red fluorescence with a maximum at 595 nm (orange-red) [28]. The green fluorescent JC-1 dye forms red fluorescent aggregates when concentrated in energized mitochondria in response to their higher membrane potential, which is affected by oxidative stress.

As displayed in Figure 5a and Figure 5b, JC-1 red/green fluorescent signal significantly decreased following cell exposure to H_2_O_2_, while treatment of cells with IntegroPectin significantly reversed this effect.

**Figure 5.**
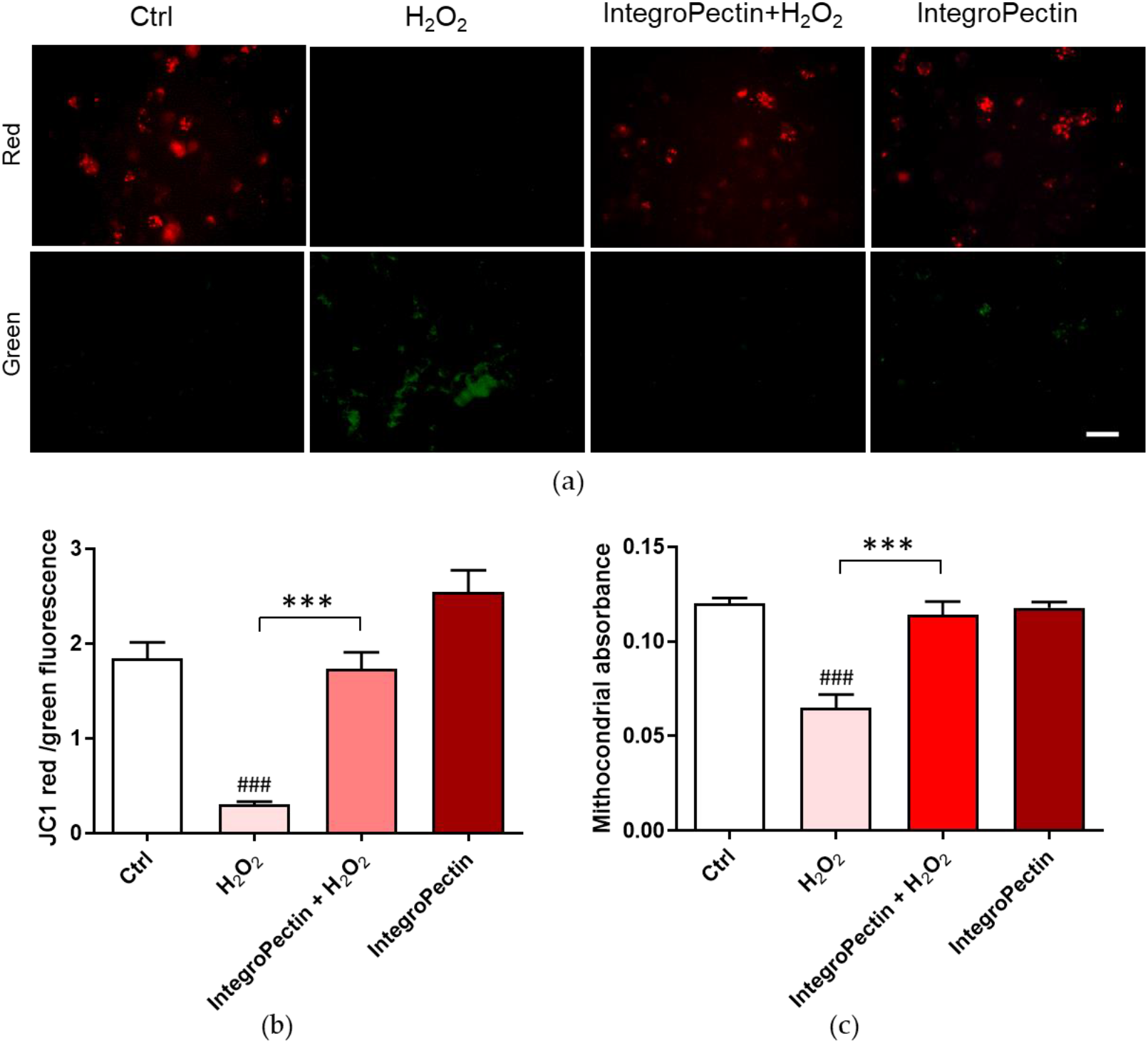
Effects of grapefruit IntegroPectin on mitochondria protection. (a) Fluorescence microscope inspection of untreated cells (Ctrl) or treated with IntegroPectin or with H_2_O_2_ alone or in combination with IntegroPectin submitted to JC-1 assay; (b) Histogram of the ratio between JC-1 red and green fluorescence intensity; (C) Histogram of mitochondria absorbance. Scale bar 100 μm. Tukey test: ### p < 0.001 as compared to control (Ctrl) group; *** p < 0.001.

Sustained mitochondrial damage triggers mitochondrial swelling due to increased colloidal osmotic pressure in the matrix accompanied by mitochondrial membrane depolarization and ATP hydrolysis. We assessed mitochondrial swelling by monitoring the decrease in light scattering at 540 nm. Treatment with the newly obtained IntegroPectin fully counteracted the significant H_2_O_2_-driven mitochondrial swelling (Figure 5c), supporting its mitoprotective function.

## 3. Discussion

Along with high *in vitro* neuroprotective and antioxidant action on human neuroblastoma cell line SH-SY5Y, grapefruit IntegroPectin extracted via hydrodynamic cavitation exerts powerful antiproliferative activity. On one hand treatment with grapefruit IntegroPectin (10 mg/ml) produces cell death. On the other a lower dose (1 mg/ml) is associated to cell cycle arrest exactly at the G_2_/M phase. Future studies will investigate whether the cell cycle arrest and antiproliferative effects observed might be indicative of cell differentiation process triggering. Undifferentiated SH-SY5Y cells, though sharing few properties with mature neurons, after differentiation to enhance their usefulness as neuronal models, have even increased oxidative vulnerability [29]. In this respect, the high tolerance to oxidation by concentrated (0.2 M) H_2_O_2_, exhibited by the SH-SY5Y cells treated with 1 mg/ml of grapefruit IntegroPectin along with the antiproliferative activity is highly promising in light of *in vivo* experiments and future practical use of this new citrus pectin as antioxidant and antitumor agent.

From a structural viewpoint, grapefruit IntegroPectin is very different when compared to commercial citrus pectin extracted via conventional hydrolytic extraction in hot acidic water followed by precipitation with alcohol. Figure 6 shows the X-ray diffraction (XRD) patterns of IntegroPectin and commercial citrus pectin.

**Figure 6.**
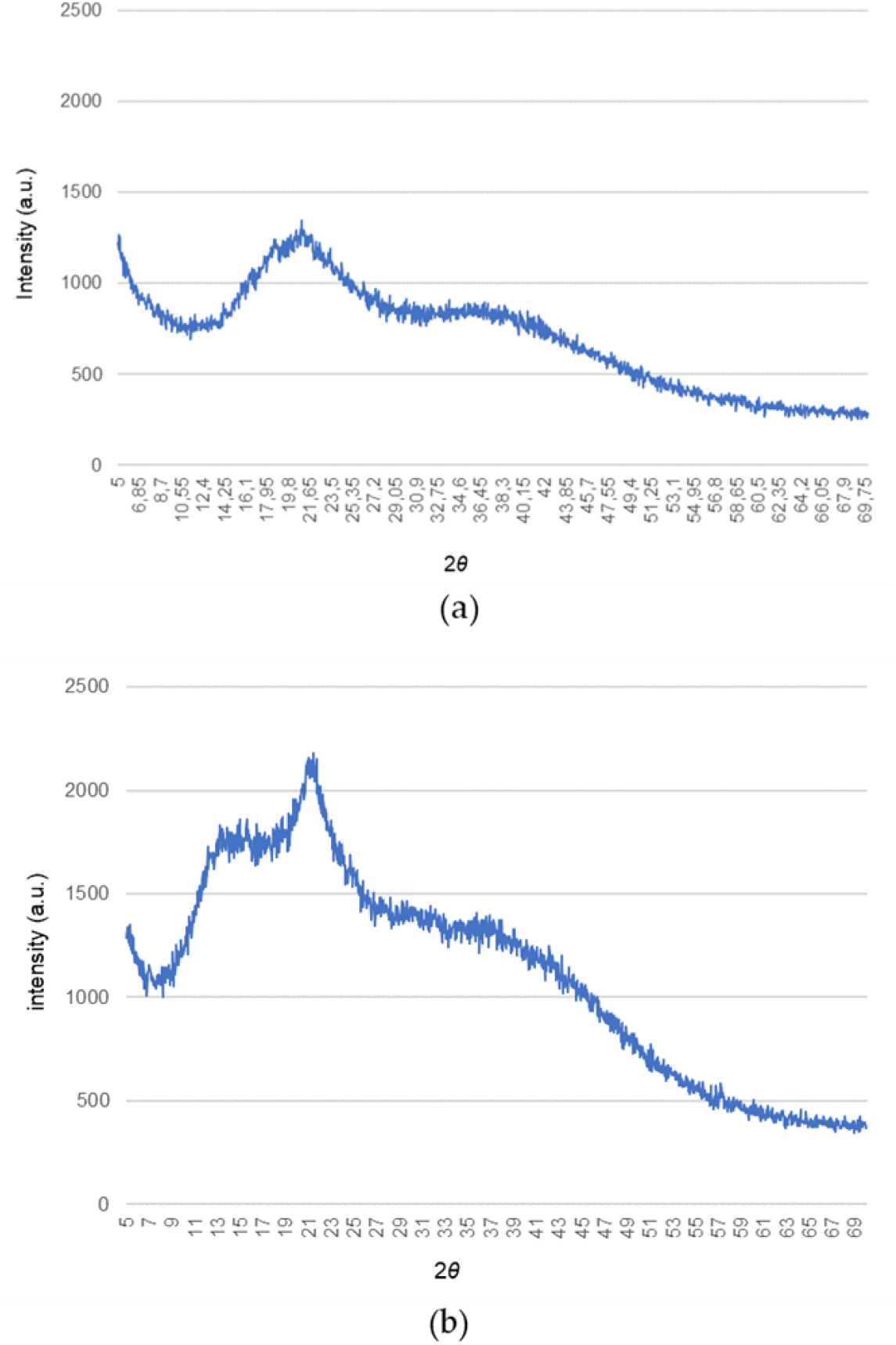
X-ray diffractograms of grapefruit IntegroPectin (a) and commercial citrus pectin (b).

Exactly as it happens with grapefruit pectin obtained via cavitation driven by ultrasounds, the diffraction peaks characteristic of commercial citrus pectin at 12.4°, 14.3°, 21.0°, 28.9°, 31.5°, 32.2°, 40.2° [30] shift to higher 2*θ* values with several sharp peaks disappearing [31].

This indicates that hydrodynamic cavitation of grapefruit biowaste induces nearly complete decrystallization of the homogalacturonan (HG) chains of grapefruit pectin following which, we remind, crystallize in hexagonal closest packing arrangement which ensures the closest packing of the chains (we also remind that diffraction from the HG regions is the only one contributing to the XRD pattern) [32]. In further detail, these findings confirm that cavitation, no matter if acoustic or hydrodynamic, destroys the “fringed-micellar” structure of the crystalline regions of the semicrystalline pectin biopolymer [33]. This finding, along with the substantially higher amount of RG-I regions and low degree of esterification (DE), explains also the significantly larger solubility of the IntegroPectin in water at room temperature when compared to the poorly soluble commercial citrus pectin. Indeed, grapefruit IntegroPectin obtained upon freeze-drying is a low-methoxyl citrus pectin (DE = 14% [23] compared to 69% for commercial citrus pectin [24]), containing also a uniquely high amount of naringin (4′,5,7-trihydroxyflavonone-7-rhamnoglucoside), approaching 74 mg/g [20], adsorbed at its surface. For comparison, the highest yield values reported for naringin extracted from fresh grapefruit albedo is 10.5 mg/g [34]. Due to its anti-cancer, anti-apoptotic, anti-atherogenic, anti-inflammatory and antioxidant properties, naringin is currently intensively researched in light of utilization for cancer prevention and treatment [35]. One of the main limitations to its use as therapeutic agent is the poor solubility in water (0.5 g/L [36]) which leads to poor pharmacokinetics [35]. It is likely that its dispersion on the highly soluble fibers of grapefruit IntegroPectin obtained upon freeze drying enhances its bioavailability. Grapefruit IntegroPectin, furthermore, is rich in adsorbed limonene, linalool and linalool oxide (predominantly the *cis* isomer) [21]. Both linalool [37] and linalool oxide [38] have been lately shown to exert neuroprotective and anticonvulsant and antinociceptive activities *in vitro* and *in vivo*, respectively.

RG-I enriched grapefruit IntegroPectin, however, does not act simply as a carrier of bioactive naringin, but possesses high bioactivity in itself which magnifies the activity of the adsorbed flavonoid. This was observed also for ultrasonically-obtained grapefruit pectin which at 2 mg/mL concentration has a free radical scavenging activity nearly twice higher than that of commercial citrus pectin [31], ascribed by Liu and co-workers to its lower viscosity in solution which would enhance effective interaction between the pectin’s hydroxyl groups and free radicals [39]. Remarkably, enzymatically extracted apple pectin enriched in galactose, rhamnose and phenolic compounds as apple pectin extracted from dried apple pomace using sulfuric acid in hot water, was recently shown to possess anti-oxidant and antitumor activity on human adenocarcinoma and melanoma cell lines [40]. Finally, in light of forthcoming *in vivo* and clinical trials of the newly developed citrus IntegroPectin it is also relevant that both pectin [41] and citrus flavonoids [42] and essential oils (terpenes and other oxygenated compounds) [43] share an excellent health and safety profile.

## 4. Materials and Methods

### 4.1. Solubilization of IntegroPectin

Grapefruit IntegroPectin was solubilized dissolving 10 mg of pectic polymer powder in 1 mL of cell culture medium. The solution was filtered using a 0.45 μm sartorius filter and stored at +4 °C.

### 4.2. Cell Cultures and Treatment

SH-SY5Y cells were cultured in T25 tissue culture flasks in Complete Dulbecco’s Modified Eagle’s Medium and F12 (DMEM/F12; 1:1), supplemented with 10% fetal bovine serum (FBS), 100 U/mL penicillin and 100 U/mL streptomycin and 2 mM L-Glutamine, in a humidified atmosphere of 95% air and 5% CO2 at 37 °C. The cell culture medium was changed every three days, and the cells were sub-cultured once they reached 90% confluence. The effects of grapefruit IntegroPectin were tested in cells cultured in 96-wells plates. All treatments were performed at least 24h after plating. Based on the experimental groups, the cells received the following treatments: H_2_O_2_ (200 μM for 24h), Integropectin (10 mg/mL, 1 mg/mL, 0.1mg/mL and 0.01 mg/mL for 24h or 48h), a combination of IntegroPectin and H_2_O_2_, with pectins administered 24h before (pretreatment) or immediately before (co-treatment). The control (Ctrl) group was treated with an equal volume of cell medium.

### 4.3. Cell Viability and Cell Morphology

Cells were grown at a density of 2 × 10^4^ cell/well on 96-well plates in a final volume of 100 μL/well. Cell viability was assessed by measuring the reduction of a yellow tetrazolium salt (3-(4,5-dimethylthiazol-2-yl)-2,5-diphenyltetrazolium bromide (MTT, 0.5 mg/mL) to purple formazan crystals by mitochondrial succinate dehydrogenase expressed in metabolically active cells after 3h incubation at 37 °C. Absorbance was measured at 570 nm with background subtraction after extracting MTT–formazan product with dimethyl sulfoxide (DMSO) 100 μL/well. Cell viability was expressed as arbitrary units, with the control group set to 1.

For the analysis of cell morphology, cells were grown at a density of 5 × 10^3^ cells/well on 96-well plates in a final volume of 100 μL/well. To this end, cells were fixed with 4% formaldehyde solution for 15 min at room temperature and washed twice with Phosphate-buffered saline (PBS). For analysis of cell nuclei morphology, nuclei were counter-stained with the fluorescent stain 4′,6-diamidino-2-phenylindole (DAPI). The cellular images obtained using the Zeiss Axio Scope 2 microscope (Carl Zeiss, Oberkochen, Germany) were analyzed with the ZEISS-ZEN imaging software, measuring each time the cell body size and the number of cell debris per field, and examining nuclei signal.

### 4.4. CellTox Green Cytotoxicity Assay

Cells were grown at a density of 2 × 10^4^ cell/well on 96-well plates in a final volume of 100 μL/well. Grapefruit Integropectin cytotoxicity was assessed by CellTox™ Green Cytotoxicity Assay (Promega Corporation, Madison, WI 53711 USA). At the end of treatment, CellTox™ Green Reagent (100μl) was added to each well. After 15 min of incubation, fluorescence was read using a Microplate Reader GloMax fluorimeter (Promega Corporation, Madison, WI 53711 USA) at the excitation wavelength of 485 nm and emission wavelength 530 nm. For positive control of toxicity, lysis solution was added to replicate wells 30 min before reading. After background subtraction, results were expressed as a percentage of the control group.

### 4.5. Flow Cytometry analysis of cell cycle

Cells were grown at a density of 1.2 × 10^5^ cell/well on 24-well plates in a final volume of 500 μL/well. At the end of treatment, cells were harvested by centrifugation, washed with PBS and incubated for 30 min in the dark in a PBS solution containing Triton X100 (0.1%), 20 μg/mL PI (Merck, Milan, Italy) and 200 μg/mL RNase (Thermo Fisher, Milan, Italy) [44]. At least 1 × 10^4^ cells were analyzed for each sample.

### 4.6. Analysis of ROS and Oxidation Kinetics

To assess intracellular ROS concentration, SH-SY5Y cells were plated at a density of 1 × 10^4^ cells/well on 96-well plates in a final volume of 100 μL/well. At the end of the treatments, 2′,7′-Dichlorofluorescin diacetate (DCFH-DA, Merck, Darmstadt, Germany, 1 mM) dissolved in PBS was added to each well and the plate was placed in the dark for 10 min at room temperature for cell uptake. Upon cleavage of the acetate groups by intracellular esterases and oxidation, the nonfluorescent DCFH-DA is converted to the highly fluorescent 2’,7’-dichlorofluorescein (DCF). The oxidation kinetics was investigated by placing SH-SY5Y cells at a density of 1 × 10^4^ cell/well on 96-well plates in a final volume of 100 μL/well. The kinetics of ROS production was evaluated for 90 minutes after the addition of H_2_O_2_. After washing with PBS, DCF fluorescence intensity was analyzed by the fluorescence Zeiss Axio Scope 2 microscope (Carl Zeiss, Oberkochen, Germany) and using a Microplate Reader GloMax fluorimeter (Promega Corporation, Madison, WI 53711 USA) at the excitation wavelength of 475 nm and emission wavelength 555 nm. Results were expressed as a percentage of the control group.

### 4.7. Mitochondrial Membrane Potential Analysis

The mitochondrial transmembrane potential was measured by incubating the cells for 30 min at 37 °C with 2 mM JC-1 red dye (5,5′,6,6′-tetrachloro-1,1′,3,3′-tetraethylbenzim-idazolylcarbocyanine iodide) using the MitoProbe JC-1 assay kit (Molecular Probes, USA). Mitochondrial depolarization is indicated by a decrease in the red/green fluorescence intensity ratio, evaluated by the aforementioned fluorimeter and fluorescence microscope equipped with a 488 nm excitation laser.

### 4.8. Swelling of Isolated Mitochondria

The mitochondrial swelling was evaluated by measuring the decrease in the absorbance of the mitochondrial suspensions. The absorbance of isolated mitochondria was monitored for 5 min at 37 °C at 540 nm, using a GloMax Discover multimode plate reader (Promega Corporation, Madison, WI 53711 USA).

### 4.9. XRD Measurements

The pectin samples were analyzed by a D5005 X-ray diffractometer (Bruker AXS, Karlsruhe, Germany) operating at 40 kV and 30 mA. The X-ray radiation was generated via a copper (Kα) anode and made monochromatic via the instrument’s secondary monochromator. The diffraction profile of both grapefruit IntegroPectin and commercial citrus pectin (galacturonic acid ≥74.0 %, dried basis) purchased from Sigma-Aldrich (Merck Life Science, Milan, Italy) at 0.15°/min acquisition rate over the 5.0°–70.0° 2*θ* range.

### 4.8. Statistical Analysis

Data analysis was performed using GraphPad Prism 5 software (GraphPad Software, San Diego, CA, USA). The results are presented as mean ± SE, and in some cases are expressed as arbitrary units, with controls equal to 1, or as percentage of control. Statistical evaluations were performed by one-way ANOVA, followed by Tukey Post-Hoc test, or t-test. Differences in *p*-value less than 0.05 were considered statistically significant.

## 5. Conclusions

Tested *in vitro* on SH-SY5Y neuroblastoma cells, grapefruit IntegroPectin rich in naringin, linalool, linalool oxide and limonene adsorbed at the outer surface was found to exert powerful neuroprotective, antioxidant and antiproliferative activites. Besides preserving the highly bioactive RG-I regions usually degraded in the conventional hydrolytic process used to produce pectin on commercial scale, the hydrodynamic cavitation extraction process nearly eliminates the crystalline regions of the semicrystalline pectic biopolymer. This ensures quick dissolution of this new pectin in water at room temperature enabling the biopolymer in solution to exert its multiple biological actions. The ability of this new pectin to inhibit cell proliferation or even kill, at high doses, neoplastic cells, coupled to the excellent health and safety profile of pectin, citrus flavonoids and essential oils, opens up new therapeutic strategies in cancer research. Indeed, cancer is a multifactorial disease, involving both endogenous/exogenous factors, in which free radicals play a key role. Cancer cells have a high free radical formation activity as compared to healthy cells, and the inhibition of this process can lead to benefits in tumor progression. It has been shown that ROS, and the subsequent oxidation of macromolecules, facilitate mutagenesis, tumor growth and metastasis [45]. Furthermore, epidemiological studies suggest that the incidence of cancer is lower in populations where the diet is rich in antioxidants such as those found in fruits and vegetables [46] Therefore, the combination of antioxidant and antiproliferative effect can represent a winning combination against tumor progression.

Similarly, the strong antioxidant properties of grapefruit IntegroPectin and its ability to preserve mitochondrial membrane potential and morphology, severely impaired in neurodegenerative disorders, make this new biomolecule an attractive therapeutic agent for oxidative stress-associated brain disorders.

Detailed molecular mechanism studies underlying the antiproliferative and neuro-protective effects of IntegroPectin are urgently needed in order to take full advantage of its vast therapeutic and preventive potential.

## Author Contributions

Conceptualization, V.D., D.N.; methodology, R.C..; software, G.M; formal analysis, C.G., M.S., P.P., A.A., S.R., F.G., A.S.; investigation, D.N., P.P., V.D., C.G., M.S.; resources, D.N., F.M., M.P, G.M.; data curation, D.N.; C.G., M.S., V.D.; writing—original draft preparation, V.D., D.N., M.P.; writing—review and editing, V.D., D.N., M.P., G.M.; supervision, V.D., D.N. All authors have read and agreed to the published version of the manuscript.

## Funding

This research received no external funding

## Data Availability Statement

All experimental data are available by contacting the corresponding Authors.

## Acknowledgments

We thank OPAC Campisi Società Cooperativa Agricola (Siracusa, Italy) for a generous gift of waste grapefruit peel from which the IntegroPectin was extracted. S.R. is research fellow funded by European Union-FESR FSE, PON Ricerca e Innovazione 2014–2020 (AIM line 1).

## Conflicts of Interest

The authors declare no conflict of interest.

## Notes

### Competing Interest Statement

The authors have declared no competing interest.

## References

1. GBD 2016 Neurology Collaborators, Global, regional, and national burden of neurological disorders, 1990-2016: A systematic analysis for the Global Burden of Disease Study 2016, Lancet Neurol. 2019, 18, 459–480. doi:10.1016/s1474-4422(18)30499-x

2. Picone, P.; Di Carlo, M.; Nuzzo, D. Obesity and Alzheimer’s disease: molecular bases, Eur. J. Neurosci. 2020, 52, 3944–3950. doi:10.1111/ejn.14758

3. Nuzzo, D.; Picone, P. Potential neurological effects of severe COVID-19 infection, Neurosci. Res. 2020, 158, 1–5. doi:10.1016/j.neures.2020.06.009

4. Picone, P.; Nuzzo, D.; Caruana, L.; Scafidi, V.; Di Carlo, M. Mitochondrial dysfunction: different routes to Alzheimer’s disease therapy, Oxid. Med. Cell. Longev. 2014, 2014, 780179. doi:10.1155/2014/780179

5. Picone, P.; Nuzzo, D. Giacomazza, D.; Di Carlo, M. β-Amyloid peptide: The cell compartment multi-faceted interaction in Alzheimer’s disease, Neurotox. Res. 2020, 37, 250–263. doi:10.1007/s12640-019-00116-9

6. Nuzzo D, Presti G, Picone P, Galizzi G, Gulotta E, Giuliano S, Mannino C, Gambino V, Scoglio S, Di Carlo M., Effects of *Aphanizomenon flos-aquae* (Klamin) extract on a cell model of neurodegeneration, Oxid Med Cell Longev. 2018, 2018, 9089016. doi:10.1155/2018/9089016

7. Nuzzo D, Contardi M, Kossyvaki D, Picone P, Cristaldi L, Galizzi G, Bosco G, Scoglio S, Athanassiou A, Di Carlo M., Heat-resistant *Aphanizomenon flos-aquae* (AFA) extract (Klamin) as a functional ingredient in food strategy for prevention of oxidative stress, Oxid. Med. Cell. Longev. 2019, 2019, 9481390. doi:10.1155/2019/9481390

8. R. Kahl, H. Kappus, Toxicology of the synthetic antioxidants BHA and BHT in comparison with the natural antioxidant vitamin E, Z. Lebensm. Unters. Forsch. 1993, 196, 329–338. doi:10.1007/bf01197931

9. Nuzzo D. Role of Natural Antioxidants on Neuroprotection and Neuroinflammation, Antioxidants 2021, 10, 608. doi:10.3390/antiox10040608

10. O. Zaitseva, A. Khudyakov, M. Sergushkina, O. Solomina, T. Polezhaeva, Pectins as a universal medicine, Fitoterapia 2020, 146, 104676. doi:10.1016/j.fitote.2020.104676

11. Daher Firas B., Braybrook S.A., How to let go: pectin and plant cell adhesion, Front. Plant Sci. 2015, 6, 523. doi:10.3389/fpls.2015.00523

12. Beukema, M., Faas, M.M., de Vos, P. The effects of different dietary fiber pectin structures on the gastrointestinal immune barrier: impact via gut microbiota and direct effects on immune cells, Exp. Mol. Med. 2020, 52, 1364–1376. doi:10.1038/s12276-020-0449-2

13. D. Seisun, N. Zalesny, Strides in food texture and hydrocolloids, Food Hydrocoll. 2021, 117, 106575. doi:10.1016/j.foodhyd.2020.106575

14. Ciriminna, R.; Chavarría-Hernández, N.; Hernández, A.R.; Pagliaro, M. Pectin: A new perspective from the biorefinery standpoint, Biofuel. Bioprod. Bioref. 2015, 9, 368–377. doi:10.1002/bbb.1551

15. R. Ciriminna, A. Fidalgo, R. Delisi, L. M. Ilharco, M. Pagliaro, Pectin production and global market, Agro Food Ind. Hi Tech 2016, 27, 17–20.

16. G. Mao, D. Wu, C. Wei, W. Tao, X. Ye, R. J. Linhardt, C. Orfila, S. Chen, Reconsidering conventional and innovative methods for pectin extraction from fruit and vegetable waste: Targeting rhamnogalacturonan I, Tr. Food Sci. Technol. 2019, 94, 65–78. doi:10.1016/j.tifs.2019.11.001

17. T. Zhang, Y. Lan, Y. Zheng, F. Liu, D. Zhao, K.H. Mayo, Y. Zhou, G. Tai, Identification of the bioactive components from pH-modified citrus pectin and their inhibitory effects on galectin-3 function, Food Hydrocoll. 2016, 58, 113–119. doi:10.1016/j.foodhyd.2016.02.020

18. F. Meneguzzo, C. Brunetti, A. Fidalgo, R. Ciriminna, R. Delisi, L. Albanese, F. Zabini, A. Gori, L. B. dos Santos Nascimento, A. De Carlo, F. Ferrini, L. M. Ilharco, M. Pagliaro, Real-scale integral valorization of waste orange peel via hydrodynamic cavitation, Processes 2019, 7, 581. doi:10.3390/pr7090581

19. D. Nuzzo, L. Cristaldi, M. Sciortino, L. Albanese, A. Scurria, F. Zabini, C. Lino, M. Pagliaro, F. Meneguzzo, M. Di Carlo, R. Ciriminna, Exceptional antioxidant, non-cytotoxic activity of integral lemon pectin from hydrodynamic cavitation, Chemistry-Select 2020, 5, 5066–5071. doi:10.1002/slct.202000375

20. A. Scurria, M. Sciortino, L. Albanese, D. Nuzzo, F. Zabini, F. Meneguzzo, R. V. Alduina, A. Presentato, M. Pagliaro, G. Avellone, R. Ciriminna, Flavonoids in lemon and grapefruit IntegroPectin, Preprints 2021, 2021020620. doi:10.20944/pre-prints202102.0620.v1

21. A. Scurria, M. Sciortino, A. Presentato, C. Lino, E. Piacenza, L. Albanese, F. Zabini, F. Meneguzzo, D. Nuzzo, M. Pagliaro, D. F. Chillura Martino, R. Alduina, G. Avellone, R. Ciriminna, Volatile compounds of lemon and grapefruit IntegroPectin, Molecules 2021, 26, 51. doi:10.3390/molecules26010051

22. A. Presentato, A. Scurria, L. Albanese, C. Lino, M. Sciortino, M. Pagliaro, F. Zabini, F. Meneguzzo, R. Alduina, D. Nuzzo, R. Ciriminna, Superior antibacterial activity of integral lemon pectin from hydrodynamic cavitation, ChemistryOpen 2020, 9, 628–630. doi:10.1002/open.202000076

23. A. Presentato, E. Piacenza, A. Scurria, L. Albanese, F. Zabini, F. Meneguzzo, D. Nuzzo, M. Pagliaro, D. Chillura Martino, R. Alduina, R. Ciriminna, A new water-soluble bactericidal agent for the treatment of infections caused by Gram-positive and Gram-negative bacterial strains, Antibiotics 2020, 9, 586. doi:10.3390/antibiotics9090586

24. Nuzzo D., Picone P., Giardina C., Scordino M., Mudò G., Pagliaro M., Scurria A., Meneguzzo F., Ilharco L.M., Fidalgo A., Alduina R., Presentato A., Ciriminna R., Di Liberto V.. New neuroprotective effect of lemon IntegroPectin on neuronal cellular model, Antioxidants 2021, 10, 669. doi: 10.3390/antiox10050669

25. Lezaja A., Altmeyer M., Inherited DNA lesions determine G1 duration in the next cell cycle, Cell Cycle 2018, 17, 24–32. doi:10.1080/15384101.2017.1383578

26. G. K. Schwartz, M. A. Shah, Targeting the cell cycle: a new approach to cancer therapy, J. Clin. Oncol. 2005, 23, 9408–9421. doi:10.1200/jco.2005.01.5594

27. B. Kalyanaraman, V. Darley-Usmar, K. J.A. Davies, P. A. Dennery, H. J. Forman, M. B. Grisham, G. E. Mann, K. Moore, L. J. Roberts, II, H. Ischiropoulos, Measuring reactive oxygen and nitrogen species with fluorescent probes: challenges and limitations, Free Radic Biol Med. 2012, 52, 1–6. doi:10.1016/j.freeradbiomed.2011.09.030

28. F. Sivandzade, A. Bhalerao, L. Cucullo, Analysis of the mitochondrial membrane potential using the cationic JC-1 dye as a sensitive fluorescent probe, Bio Protoc. 2019, 9, e3128. doi:10.21769/BioProtoc.3128

29. J. I. Forster, S. Köglsberger, C. Trefois, O. Boyd, A. S. Baumuratov, L. Buck, R. Balling, P. M. A. Antony, Characterization of differentiated SH-SY5Y as neuronal screening model reveals increased oxidative vulnerability, J. Biomol. Screen. 2016, 21, 496–509. doi:10.1177/1087057115625190

30. R. Sharma, S. Kamboj, R. Khurana, G. Singh, V. Rana, Physicochemical and functional performance of pectin extracted by QbD approach from *Tamarindus indica* L. pulp, Carbohydr. Polym. 2015, 134, 364–374. doi:10.1016/j.carbpol.2015.07.073

31. W. Wang, X. Ma, P. Jiang, L. Hu, Z. Zhi, J. Chen, T. Ding, X. Ye, D. Liu, Characterization of pectin from grapefruit peel: A comparison of ultrasound-assisted and conventional heating extractions, Food Hydrocoll. 2016, 61, 730–739 doi:10.1016/j.foodhyd.2016.06.019

32. K.J. Palmer, M.B. Hartzog, An X-ray diffraction investigation of sodium pectate, J. Am. Chem. Soc. 1945, 67, 2122–2127. doi:10.1021/ja01228a022

33. R.M. Gohil, Synergistic blends of natural polymers, pectin and sodium alginate, J. Appl. Polym. Sci. 2010, 120, 2324–2336. doi:10.1002/app.33422

34. M. M. Victor, J. M. David, M. C.K. Sakukuma, E. L. França, P. V. J. Nunes, A simple and efficient process for the extraction of naringin from grapefruit peel waste, Green Process. Synth. 2018, 7, 524–529. doi:10.1515/gps-2017-0112

35. Ghanbari-Movahed M., Jackson G., Farzaei M. H., Bishayee A., A systematic review of the preventive and therapeutic effects of naringin against human malignancies, Front. Pharmacol. 2021, 12, 639840. doi:10.3389/fphar.2021.639840

36. G. N. Pulley, Solubility of naringin in water, Ind. Eng. Chem. Anal. Ed. 1936, 8, 360. doi: 10.1021/ac50103a020

37. R. Migheli, G. Lostia, G. Galleri, G. Rocchitta, P. A. Serra, V. Bassareo, E. Acquas, A. T. Peana, Neuroprotective effect of (R)-(−)-linalool on oxidative stress in PC12 cells, Phytomedicine Plus 2021, 1, 100073. doi:10.1016/j.phyplu.2021.100073

38. F. Negromonte Souto-Maior, D. Vilar da Fonsêca, P. R. Rodrigues Salgado, L. de Oliveira Monte, D. Pergentino de Sousa, R. Nóbrega de Almeida, Antinociceptive and anticonvulsant effects of the monoterpene linalool oxide, Pharm. Biol. 2017, 55, 63–67. doi: 10.1080/13880209.2016.1228682

39. J. Ro, Y. Kim, H. Kim, S.B. Jang, H.J. Lee, S. Chakma, J.H. Jeong, J. Lee, Anti-oxidative activity of pectin and its stabilizing effect on retinyl palmitate, Korean J. Physiol. Pharmacol. 2013, 17, 197–201. doi:10.4196/kjpp.2013.17.3.197

40. A. Wikiera, M. Grabacka, L. Byczyński, B. Stodolak, M. Mika, Enzymatically extracted apple pectin possesses antioxidant and antitumor activity, Molecules 2021, 26, 1434. doi:10.3390/molecules26051434

41. EFSA Panel on Food Additives and Nutrient Sources added to Food), Scientific Opinion on the re-evaluation of pectin (E 440i) and amidated pectin (E 440ii) as food additives, EFSA J. 2017, 15, 4866. doi:10.2903/j.efsa.2017.4866

42. O.M. Ahmed, S.F. AbouZid, N.A. Ahmed, M.Y. Zaky, H. Liu, An up-to-date review on citrus flavonoids: chemistry and benefits in health and diseases, Curr. Pharm. Des. 2021, 27, 513–530. doi:10.2174/1381612826666201127122313

43. H. Bora, M. Kamle, D.K. Mahato, P. Tiwari, P. Kumar, *Citrus* essential oils (CEOs) and their applications in food: an overview, Plants 2020, 9, 357. doi:10.3390/plants9030357

44. I. Restivo, L. Tesoriere, A. Frazzitta, M.A. Livrea, A. Attanzio, M. Allegra, Anti-poliferative activity of a hydrophilic extract of manna from *Fraxinus angustifolia* Vahl through mitochondrial pathway-mediated apoptosis and cell cycle arrest in human colon cancer cells, Molecules 2020, 25, 5055. doi: 10.3390/molecules25215055

45. Perillo, B.; Di Donato, M.; Pezone, A.; Di Zazzo, E.; Giovannelli, P.; Galasso, G.; Castoria, G.; Migliaccio, A., ROS in cancer therapy: the bright side of the moon, Exp. Mol. Med. 2020, 52, 192–203. doi:10.1038/s12276-020-0384-2

46. C. Esquivel-Chirino, J. Esquivel-Soto, J.A. Morales-González, D. Montes Sánchez, J.L. Ventura-Gallegos, L.E. Hernández-Mora, A. Zentella-Dehesa, Inflammatory Environmental, Oxidative Stress in Tumoral Progression In Oxidative Stress and Chronic Degenerative Diseases - A Role for Antioxidants, IntechOpen, Zagreb: 2013; pp.187–208. doi:10.5772/51789

